# Encoding of T=1 virus capsid structures through the interfaces of oligomer subcomponents

**DOI:** 10.1101/2024.06.27.600969

**Authors:** Mads Jeppesen, Ingemar André

## Abstract

Virus capsid formation is one the most complex self-assembly processes in nature, involving the association of a large number of protein subunits into well-defined structural states. The ability of protein subunits to efficiently self-assembly is encoded in the protein sequence, and ultimately in the protein-protein interfaces within the growing capsid. The relative strengths of interfaces can be important in directing the self-assembly process, and determine which intermediates are formed early in the self-assembly process. In this study we characterize the interfaces in homomeric T=1 virus capsids to investigate to what extent interfaces within the assemblies are different from other protein-protein interfaces, and which interfaces are most critical for self-assembly. Interfaces were divided into dimers, trimers, and pentamers and compared to interfaces of non-viral homomeric dimers, trimers, and pentamers. The analysis suggests that viral interfaces are larger than non-viral counterparts, and differ in amino acid content, but are energetically similar in terms of the quality of intermolecular interactions. Trimers are predicted to be the most stable oligomers, which may imply that they form early in the self-assembly process. However, dimeric and pentameric interfaces are typically similar in terms of predicted stability suggesting that assembly formation in T=1 capsids may progress through many different routes, rather than progressing through a single dominant intermediate species. With symmetric docking calculations, the energy landscape of the assembled capsid was characterized, and the results highlight that the assemblies exhibit deeply funneled energy landscapes encoded by protein-protein interfaces that have a high degree of specificity.

## Introduction

Formation of icosahedral virus capsids is one the most complex protein self-assembling reactions in nature, involving up to several hundred contributing subunits. Efficient self-assembly requires subtle tradeoffs between interaction specificity and kinetic accessibility[1], where specificity increases the likelihood of forming productive intermediates while simultaneously reducing the rate of capsid formation. Specificity is ultimately encoded in protein interfaces; protein-protein interfaces within the protein capsid as well as interfaces formed between capsid proteins and nucleic acids. Even the simplest form of protein capsid contains several different types of protein-protein interfaces. The properties of these interfaces, including their stabilities and relative strengths have a substantial impact on the assembly mechanism and the efficiency of capsid formation. In this study, we aim to characterize protein-protein interfaces within icosahedral protein capsids from a structural and energetic standpoint to better understand how evolution has optimized protein interactions for efficient capsid assembly.

The simplest icosahedral protein capsids, with a triangulation number of 1 (T=1), containing 60 subunits (protomers) arranged in a lattice with 30 two-fold, 20 three-fold, and 12 five-fold centers of symmetry[2], see Figure 1. Larger capsids with more than 60 subunits can be formed by associating protomers into asymmetric units where each protein chain adopts a slightly different conformation (T>1). The asymmetric unit can also consist of several different types of protein chains, forming heteromeric rather than homomeric protein capsids. Quasi-equivalence (T>1) results in an expansion of the number of types of interfaces and may include pseudo-hexameric centers. Icosahedral virus capsids thus typically have at least three types of interfaces -dimers, trimers, and pentamers – but can contain over 50 unique interfaces[3].

**Figure 1:**
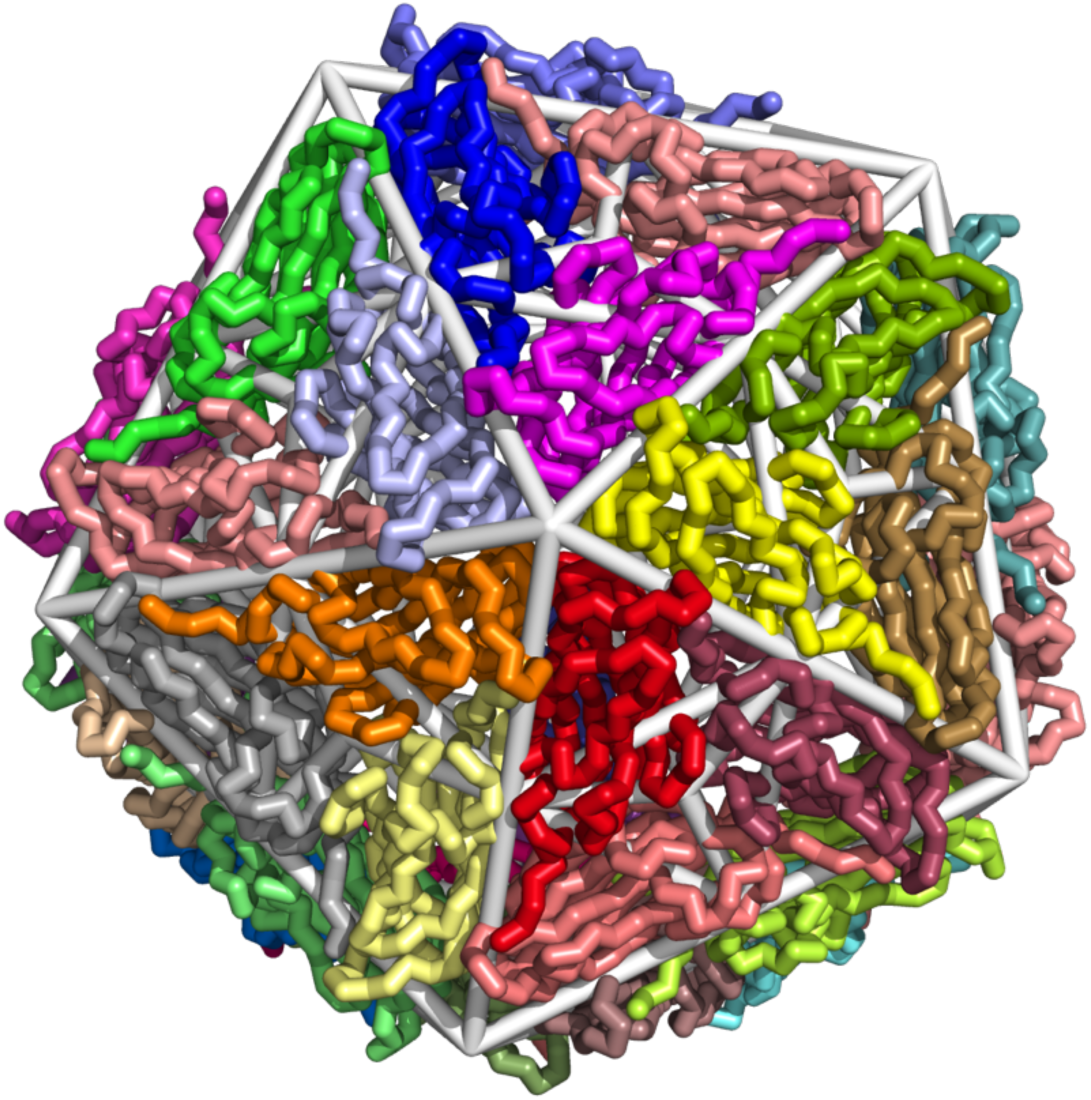
T=1 protein capsid with icosahedral symmetry. Structure of Satellite Panicum Mosaic Virus capsid[4], with icosahedral lattice shown in grey bars (taken from Viper database)[5]. The 2-fold interface between blue and light blue, 3-fold between red, raspberry, and salmon color, and 5-fold between red, orange, light-blue, magenta, and yellow subunits.

For many viruses, the capsid assembly is triggered by the binding of nucleic acids to protein subunits[6], either by increasing the effective subunit-subunit affinity or by triggering a conformation change in the bound subunit thus making them assembly competent. There are, however, many examples of virus capsid protein proteins that can assemble into capsids in the absence of nucleic acids. These systems have served as invaluable models to investigate the mechanism of virus capsid assembly, both in experiments and simulations. Experimental work has demonstrated that capsid assembly formation proceeds through the formation of a critical nucleus, followed by a growth phase where one or several subunits are added until the final capsid structure is assembled. Intermediates are often found along the assembly pathway, either early intermediates associated with the formation of the critical nucleus[7] or late intermediates associated with partially assembled capsids[8, 9].

Smaller oligomers associated with the symmetry centers in the icosahedra (dimers, trimers, and pentamers) are natural building blocks in the assembly models of icosahedral capsids, and it is typically postulated that initial assembly proceeds through these oligomers[6, 7, 9–11]. Tracking the formation of individual oligomers during assembly is difficult to achieve experimentally, but data on the overall stability of capsids indicate that protein-protein interfaces in capsids are weak (−3-4 kcal/mol)[6, 12]. Models suggest that efficient self-assembly requires weak interactions to reduce kinetic traps[6, 13, 14]. High-affinity interfaces can result in over-nucleation, with too few monomeric building blocks available to extend the nucleus[7, 15, 16]. Strong interactions can also lead to trapping of subunits into intermediate clusters with defective shells that form alternative structures or open assemblies[17, 18]. Weak interaction enables correction of the mis-associated states, leading to an efficient assembly[19]. On the other hand, strong interactions may be associated with structural defects which may be beneficial for some types of viruses[13].

The potential for kinetic trapping suggests that interface formation has been evolutionarily optimized for assembly efficiency. The relative affinity between oligomeric species will decide on the assembly pathway, as has been observed with other types of homooligomers[20]. If for example pentamers are associated with high affinity, they are expected to form rapidly leading to an assembly mechanism driven by pentamer-pentamer interactions. On the other hand, a similar stability of dimers, trimers, and pentamers would suggest a more complex assembly pathway with many more intermediate states. It is thus of interest to investigate the relative stability of oligomeric interfaces formed in icosahedral interfaces. The importance of weak interactions for efficient self-assembly may indicate that oligomers within capsids have special interfaces compared to other protein-protein interfaces. It is thus of interest to compare virus capsid oligomers with oligomers from other proteins.

Bahadur et al. analyzed the interfaces of 49 T=1, T=3, and pseudo T=3 protein virus capsids[3], characterizing interface sizes, amino acid composition, and burial of polar and apolar atoms and compared interfaces with homodimers and protein complexes. More recently Chen and Brooks[21] carried out a comparison of the structure of interfaces between capsids and regular protein interfaces using a structural alignment approach and demonstrated a statistically significant difference. In this study, we focus on fully symmetric T=1 viral capsids and compare the interfaces of dimeric, trimeric, and pentameric subcomponents to C_2_-, C_3_-, and C_5_-symmetric homomeric proteins. We compare the structure and energetics of interface formation in these sets, which can give insight into how assembly mechanisms are encoded in individual interfaces with protein capsids and how these systems have evolved. It can also increase our understanding of how assembly mechanisms may be encoded into designed proteins for efficient formation of cage structures[22].

## Results

We focus here on naturally occurring icosahedral T=1 protein cages from viral proteins. T=1 capsids have perfect symmetry and can be compared to perfectly symmetric dimers, trimers, and pentamers found in the protein data bank. The perfect symmetry also enables us to model the structure and interactions with Rosetta, which has the capabilities to model complex symmetries like icosahedra and carry out symmetric docking calculations. A non-redundant set of 45 T=1 viral capsids were harvested from the Viper database[5] (Table S1). For each capsid, we extract the major oligomeric components. 5-folds can be formed by first-neighbor chains or second-neighbor chains[3]. We focus only on nearest neighbor interfaces although the other interface is still used in energy calculations. To improve the quality of the structures, solved at resolutions up to 3.6 Å resolution (Table S1), they were subjected to symmetric all-atom refinement[23] with the Rosetta energy function[24] before structural and energetic metrics were calculated.

To understand if viral interfaces are different from other protein-protein interfaces, we collected a non-redundant set of homodimers, trimers, and pentamers from the Protein Data Bank [25]. The sets consisted of 8173, 912, and 132 dimers, trimers, and pentamers respectively. To reduce the influence of protein chain length on the results, we subsampled a set of structures from each symmetry type so that they had a similar length probability distribution. This reduced the set of dimers, trimers, and pentamers to 2693, 296, and 62 respectively. The sampled length distributions are highly similar to the length distributions for the capsid, although the C_5_ oligomers lack some structures above 500 residues (Figure S1). Because the compared data in this study are generally normally distributed, we used Wilcoxon rank-sum tests to compare data sets[26]. This test evaluates whether two sets were drawn from the same distribution, but does not directly evaluate the difference in mean values.

### Surface burial in viral interfaces compared to homomers

The buried surface area per subunit upon assembly formation was calculated and is presented in Figure 2 and the average values across all members in sets in Table 1. The buried surface area per interface is also presented in Table 1. In general, the distributions are broad for all interface types with large standard deviations. The averages for the C_3_ (I:C_3_) interfaces are substantially larger than the corresponding values for C_2_ (I:C_2_) and C_5_ (I:C_5_) within the T=1 capsids, while the I:C_5_s have slightly bigger values than the I:C_2_s. This is very different from the data presented by Bahadur et al. on all interfaces in T=1 and T=3 capsids, where dimers (2260 Å^2^) are substantially larger than trimers (1480 Å^2^) and pentamers (1980 Å^2^), with trimers being the smallest. The I:C_3_ surface size distribution has two main modes, with a set of very large interfaces dominating the average (Figure 2A). This can be observed in the difference between the mean and median values (Table 1). The opposite is true for the I:C_2_ and I:C_5_ interfaces, where the mean interface is substantially smaller than the median. The interfaces of the homomeric interfaces (C_2_, C_3_ and C_5_) are consistently smaller than the interfaces from the capsid subcomponents and while the C_3_ trimers are largest on average the capsid trimers bury double the amount of surface area in comparison. Bahadur et al[3] compare ISASA values to values from a homodimeric set of 122 complexes that was originally presented by Bahadur et al[27]. The average values are substantially larger than this set than ours, 1940 Å^2^ compared to 1365 Å^2^. We have confirmed that the structures presented in both sets have similar ISASA values, so the difference is likely due to the different set homodimers (our set containing 2693 dimers) and selection on chain length.

**Figure 2:**
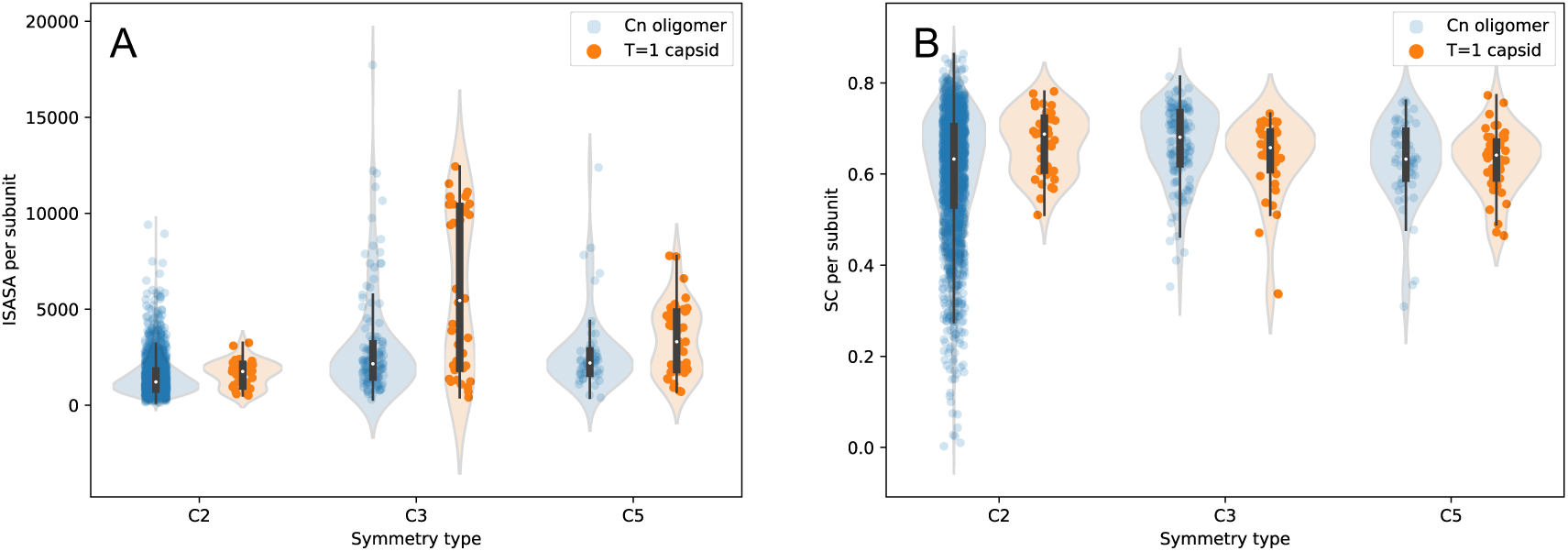
Comparisons of interfaces between viral oligomers and homomers from PDB. Comparison of symmetrical subcomponents in T=1 virus capsid proteins and C2, C3, and C5 symmetric homomers. The black box shows the quartiles, while the whiskers extend to show the limits of the distribution. The violin plot shows a kernel density estimate of the data. **A)** Buried surface area upon assembly formation (ISASA) per subunit. **B)** Shape complementarity of interfaces calculated by S_c_.

**Table 1.**

| Parameter | Capsid |  |  | Oligomers |  |  |
| --- | --- | --- | --- | --- | --- | --- |
|  | I:C<sub>2</sub> | I:C<sub>3</sub> | I:C<sub>5</sub> | C<sub>2</sub> | C<sub>3</sub> | C<sub>5</sub> |
| ISASA<sup>a</sup> (Å<sup>2</sup>) per interface (mean) | 1628 | 2995 | 1707 | 1467 | 1504 | 1356 |
| ISASA<sup>a</sup> (Å<sup>2</sup>) per interface (median) | 1757 | 2728 | 1653 | 1215 | 1079 | 1653 |
| No. contacts per subunit<sup>b</sup> (mean) | 64 | 321 | 342 | 59 | 161 | 266 |
| No. contacts per subunit<sup>b</sup> (median) | 62 | 276 | 338 | 50 | 123 | 215 |
| Shape complementarity (SC) (mean) | 0.67 | 0.63 | 0.63 | 0.60 | 0.67 | 0.63 |
| Shape complementarity (SC) (median) | 0.69 | 0.66 | 0.64 | 0.63 | 0.68 | 0.63 |
| ΔG (REU)<sup>c</sup> per subunit and ISASA per 1000Å (mean) | -21 | -32 | -25 | -20 | -25 | -32 |
| ΔG (REU)<sup>c</sup> per subunit and ISASA per 1000Å (median) | -20 | -36 | -25 | -17 | -22 | -28 |
| <sup>a</sup>ISASA: Buried Solvent Accessible Surface Area. <sup>b</sup>Number of contacts across the interface with a 8Å distance cutoff. <sup>c</sup>Binding energy estimated with the Rosetta energy function in units of the Rosetta energy function (REU). |  |  |  |  |  |  |

### Interface contacts and shape complementarity

The number of residues in the interface was also calculated and presented in Table 1. The 3- and 5-fold interfaces have substantially more residues within the vicinity of their binding partners than the 2-folds. The viral 2-fold and 3-folds have a greater number of interface residues than the corresponding C_2_ and C_3_s, in particular the viral trimers. Many interacting residues do not necessarily imply strong interactions or tightly packed interfaces. To evaluate the packing of subunits at the interfaces we evaluated the shape complementarity S_c_ metric developed by Lawrence and Colman[28] as implemented in Rosetta. Viral 2-folds are found to be better packed than the 3- and 5-fold interfaces (Figure 2A) on average, but there is no strong statistical significance to these differences (I:C_2_ vs I:C_3_ p-values 0.1, I:C_2_ vs I:C_5_ p-value 0.012). The C_2_ symmetric are less packed than the viral dimers. This is due to the tail of C_2_s that have very low S_c_ values. There is likely a significant fraction of homodimers in the PDB that are either weak complexes or whose assembly state may have been misassigned.

### Amino acid preferences in interfaces

Protein interfaces have very different amino acid composition compared to surfaces of monomeric proteins, and are enriched in hydrophobic and aromatic residues but depleted in charged residues[29]. Here we are interested in characterizing the amino acid composition of viral subcomponent interfaces and their corresponding multimers, to understand if viral interfaces are different.

Figure 3 shows the amino acid composition in the interface of viral subcomponents and the frequencies of corresponding homomers. Comparisons of sets were carried out with the Wilcoxon rank-sum test, with p-values less than 10^−3^ deemed significant. The aggregated data for all sites demonstrate the enrichment of asparagine, threonine, and proline residues with depletion of alanine, leucine, valine, and glutamic acid in viral interfaces. These trends are consistent for the 2-,3- and 5-fold viral interfaces in comparison with their corresponding oligomers. For the 2-fold, arginine has a significant enrichment over the set of C_2_ dimers. There is a depletion of lysine, which may indicate a lysine-to-arginine switch in these interfaces. Taken together, these results indicate that the viral interfaces have less hydrophobic (45% vs. 49%, p-value 9×10^−8^, Figure 3), more polar amino acids (31% vs. 22%, p-value 7×10^−32^), and less charged (24% vs. 29%, p-value 5×10^−11^) than their homomeric counterparts. The enrichment of prolines in viral interfaces may be due to the constraint on the backbone of viral proteins to fold into structures in highly packed environments.

**Figure 3:**
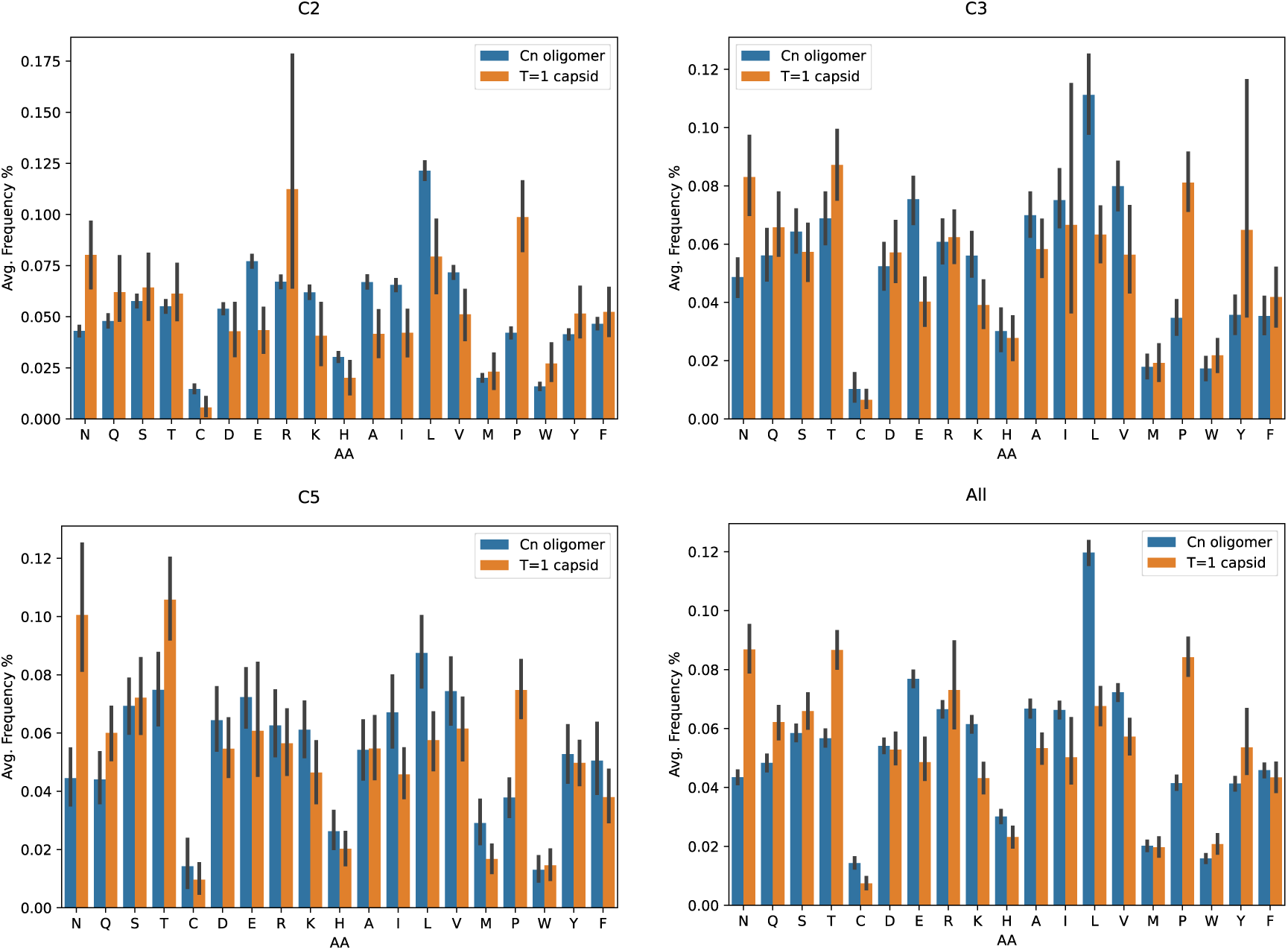
Amino acid composition in interfaces of viruses and homooligomers. Comparison of 2-, 3- and 5-fold sites in virus capsids and C_2_, C_3_, and C_5_ symmetric homomers, as well as all interface residues. Error bars correspond to 95% confidence intervals. Amino acids are ordered into the groups hydrophilic (N,Q,S,T,C), charged (D,E,R,K,H) and hydrophobic (I,L,V,M,P,W,Y,F). Since the interface analysis involves the utilization of CB atoms, glycine is not included in the analysis.

### Stability of protein interfaces

The stability of a protein interface is in not generally correlated with the size of an interface, although some weak correlations have been observed with buried surface area and stability for complexes where the bound and unbound state have less than 1Å RMSD difference[30]. Accurate predicting the ΔG for association of protein chains is an unsolved problem in computational chemistry, while substantial correlations are observed for the effect of mutations with structure-based ΔΔG calculations[31]. ΔΔG prediction methods using the Rosetta energy function[24] is one of the most successful methodologies for structure-based stability predictions[32]. While absolute ΔG predictions are not reliable with an energy-based method such as Rosetta, it can be used to compare and rank models in protein-protein docking calculations[33–35] and protein design[36]. We have also demonstrated that it can be used to predict oligomeric states of coiled-coils[37]. By comparing computed stabilities between oligomeric subcomponents with the same basic structure of the chain, the problem becomes a problem of ranking models, which is more feasible than absolute ΔG calculations. We can also compare distributions of values computed across a large set of oligomers (viral oligomers vs. homomers), which is more reliable than individual comparisons. Based on these considerations we predicted the binding energy of association by calculating the difference between the Rosetta energy of symmetrically energy-refined oligomers and the same system in an unassociated state but with repacking of sidechains. We have shown previously that ΔG computed with Rosetta normalized per subunit is a good predictor of the relative stability of oligomers[37]. These ΔG values are not on the kJ/mol scale and scale with the number of atoms that are present in the interface. This size dependence can be reduced by normalizing the ΔG per subunit by the buried surface area. The results are shown in Figure 5, with averages and medians in Table 1. We have filtered the data to remove the data points with ΔG above 0 because these are complexes for which the energy refinement simulation failed.

There is a wide distribution of stability values for each oligomerization state, with the largest range for the trimers (Figure 4A). The mean normalized stability values suggest that viral trimers are the most stable of the viral oligomers, but there is not very strong statistical support for this (I:C_3_ vs I:C_2_, p=0.01, I:C_3_ vs I:C_2_ p=0.14). Overall, the viral oligomers have similar computed stability values compared to the C_2_, C_3,_ and C_5_ sets, except for the C_3_ set which has lower mean computed stabilities than its corresponding viral trimers (I:C_3_). However, because if the large spread, there is no statistically significant difference between the data sets (p-value p=0.11). With the energy function, we can evaluate the contribution of different intermolecular interaction energy terms to the stability of the subcomponents. The van der Waals energy, hydrogen bonding, and solvation energy (modeling polar and apolar solvation) between bound and unbound states are shown in Figure 4. The result for the van der Waals energy shows that the I:C_2_ and I:C_5_ have substantially fewer packing interactions than the corresponding symmetric oligomers (p=3×10^−36^ and 3×10^−9^, respectively). The viral dimers and pentamers have better solvation energy than the set of C_2_ and C_5_ (p=1×10^−20^ and 4×10^−6^), typically suggesting more hydrophobic burial and/or less buried polar residues. This can appear incongruent with the depletion of hydrophobic residues in the viral interfaces. However, the simple frequency picture does not account for the quality of interactions. There is no substantial difference in the formation of hydrogen bonds between these sets.

**Figure 4:**
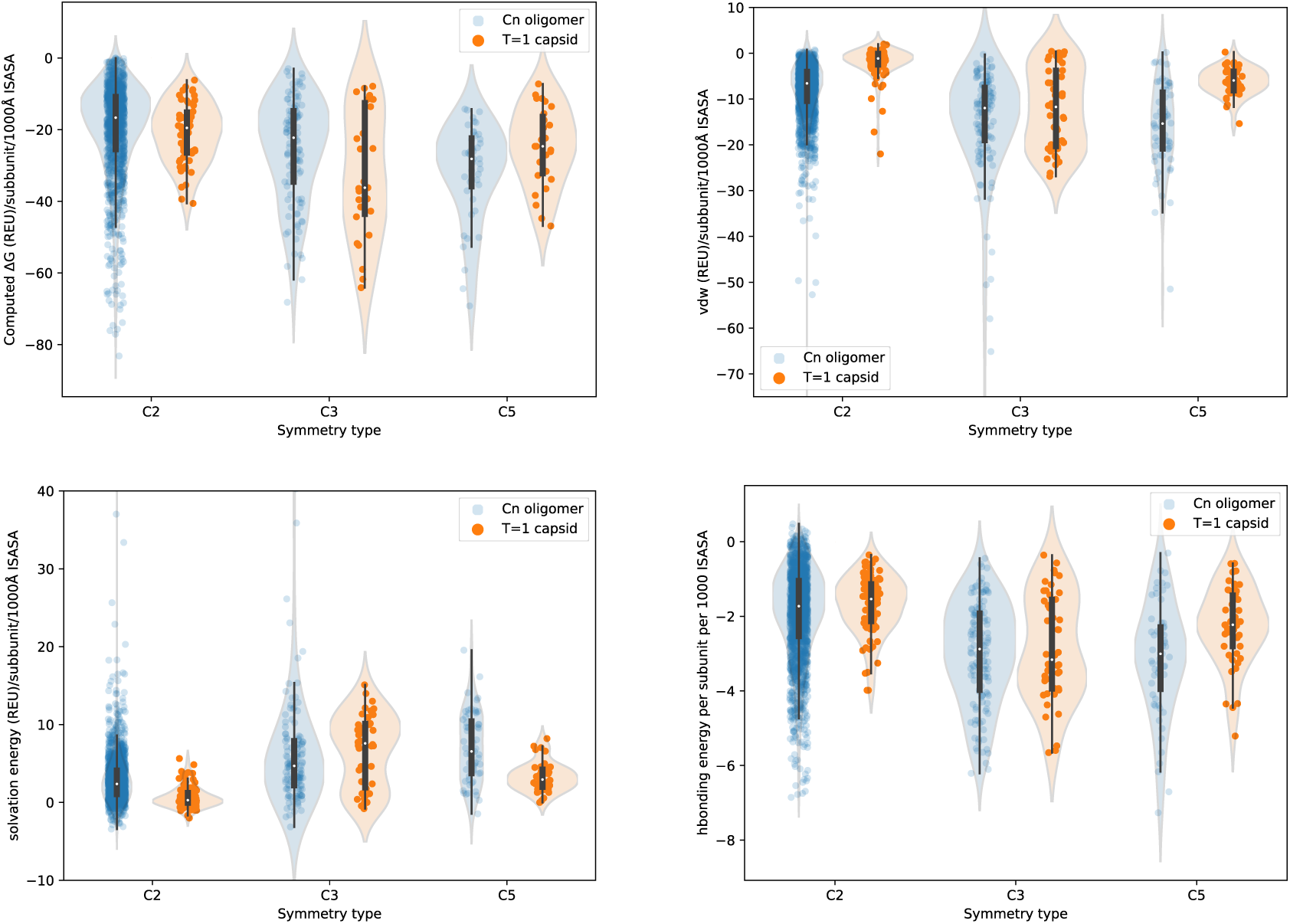
Comparison of the energetics of interface formation. Comparison of viral subcomponents with 2-, 3- and 5-fold symmetry and C_2_, C_3,_ and C_5_ symmetric oligomers normalized per subunit and surface area. Top left, computed ΔG. Top right, Van der Waals energy (attractive – repulsive Lennard-Jones in the Rosetta energy function). Bottom left, solvation energy (fa_sol term in the Rosetta energy function). Bottom right, hydrogen bonding energy (the aggregated hydrogen bonding energy terms in the Rosetta energy function).

### Energy landscapes of T=1 capsids

To evaluate how optimized the T=1 protein capsids are towards binding the assembled state found in the experimentally solved structure, we carried out symmetric docking calculations to map out the energy landscape. Symmetric EvoDOCK[35] enables local docking calculations using icosahedral symmetry to map out energy landscapes using the all-atom Rosetta energy function. We selected 20 T=1 capsids from our set for more detailed analysis. Based on the positions of subunits in the experimental structures the rigid body positions were perturbed. This perturbation was done by randomly varying the six rigid body parameters used to model the icosahedral symmetry of the system. Symmetric EvoDOCK then simulates the assembly process by minimizing the Rosetta energy function in an optimization algorithm that varies sidechain conformations and rigid body parameters. The process is repeated in a stochastic manner and the result is an energy landscape in which the interface energy (calculated ΔG but without repacking of sidechains in the unbound state) is plotted against RMSD towards the experimental structure. The results are shown in Figure 5. In all but three cases (PDB ID: 1DNV, 8ER8 and 4MGU) the minimum is found near the experimental structure (<2 Å for the symmetric subsystem used for RMSD calculations). The three failures were associated with issues with conformational sampling for systems with long loop segments. For some systems (PDB ID: 7EPP, 7TJE, and 7NO0) there are low energy minima in the neighborhood of the native state, corresponding to alternative assembly states that could be sampled during assembly. Nonetheless, in general, the energy landscape is very funneled with one deep minimum near the experimental assembly structure. This suggests that capsids explore only capsid geometries that are near the experimentally determined structure. A caveat of the analysis is the lack of backbone flexibility and imposition of strict icosahedral symmetry, which limits the conformational states that can be sampled in the simulation.

**Figure 5:**
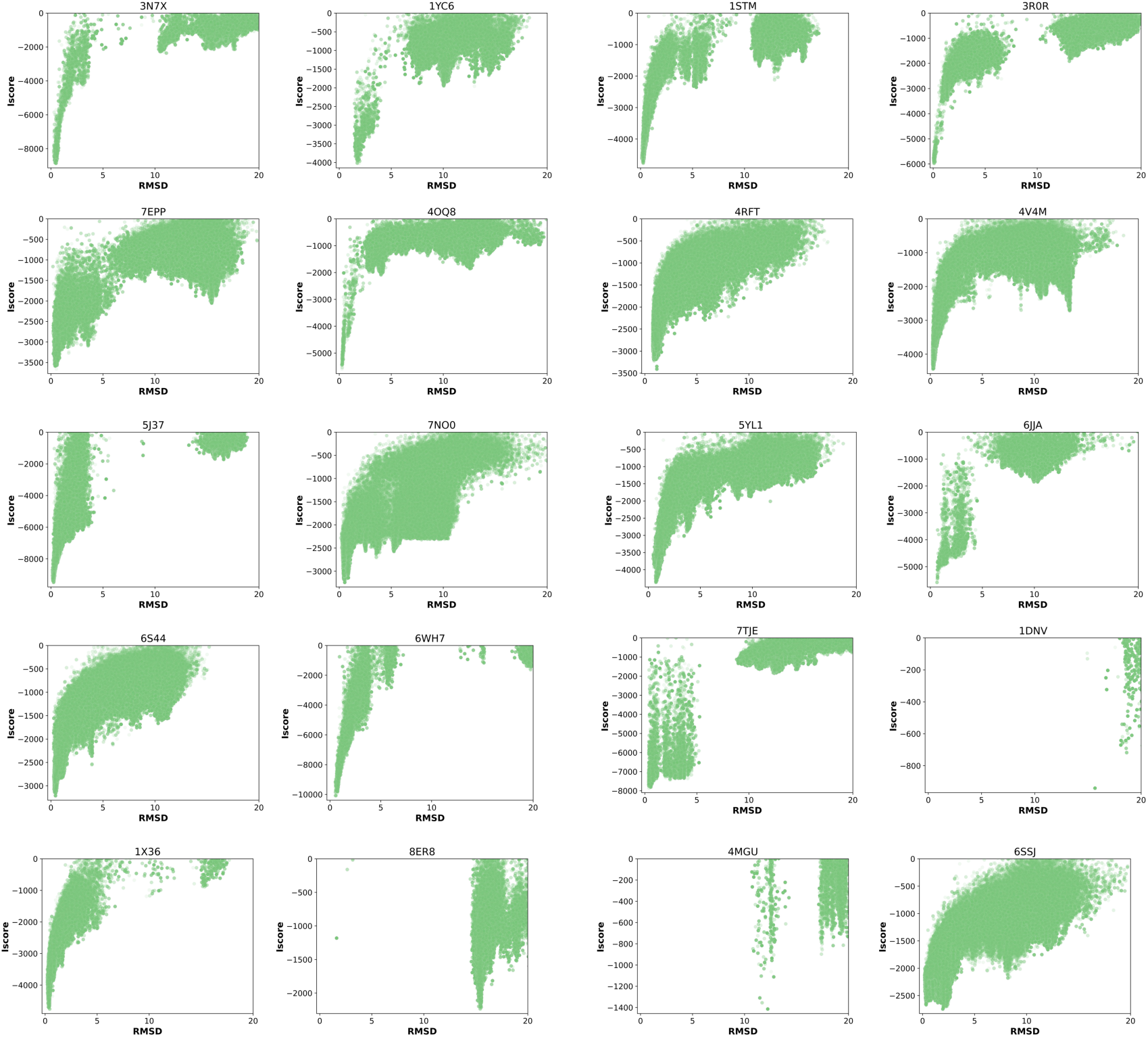
Energy landscape of T=1 capsids sampled by local symmetric docking. The interface energy (Iscore) vs. the rmsd to the native experimental structure is plotted in a scatter plot, where each dot is a model generated by the docking program.

The very lowest energy models represent an ensemble of conformations that could potentially be sampled in the assembled state. Is the variation in capsid geometry primarily associated with one type of interface that is more malleable than the others? Are the more malleable interfaces associated with the least stable oligomers in the assembly? To address these questions, we analyzed the capsid models with the lowest 2% interface energy. Each model was separated into dimers, trimers, and pentamers and aligned with the corresponding oligomer in the experimental structure to evaluate the root mean square deviation (RMSD). To facilitate comparison, we align two consecutive subunits in the oligomers to remove the size dependence of the RMSD value. The mean RMSD values for the low-energy dimers, trimers, and pentamers were calculated for each capsid, the results are shown in Figure 6A. Overall the RMSD distributions for dimers, trimers, and pentamers are highly similar. There are, however, some statistically significant differences in the mean RMSD between dimers, trimers and pentamer. In Figure 6B we rank the mean RMSD values of the oligomers in terms of magnitude and count the occurrences of different ranks. In all cases are the differences statistically significant (p-values < 10^−5^ between the first and second largest mean RMSD values), but the magnitude of the differences are small. The most common rank is to have the largest RMSD values in the pentamer followed by the dimer and trimer (5:3:2). This data suggests that pentameric interfaces are often the most malleable in terms of alternative packing orientations. Nonetheless, there does not appear to be a strong preference.

**Figure 6:**
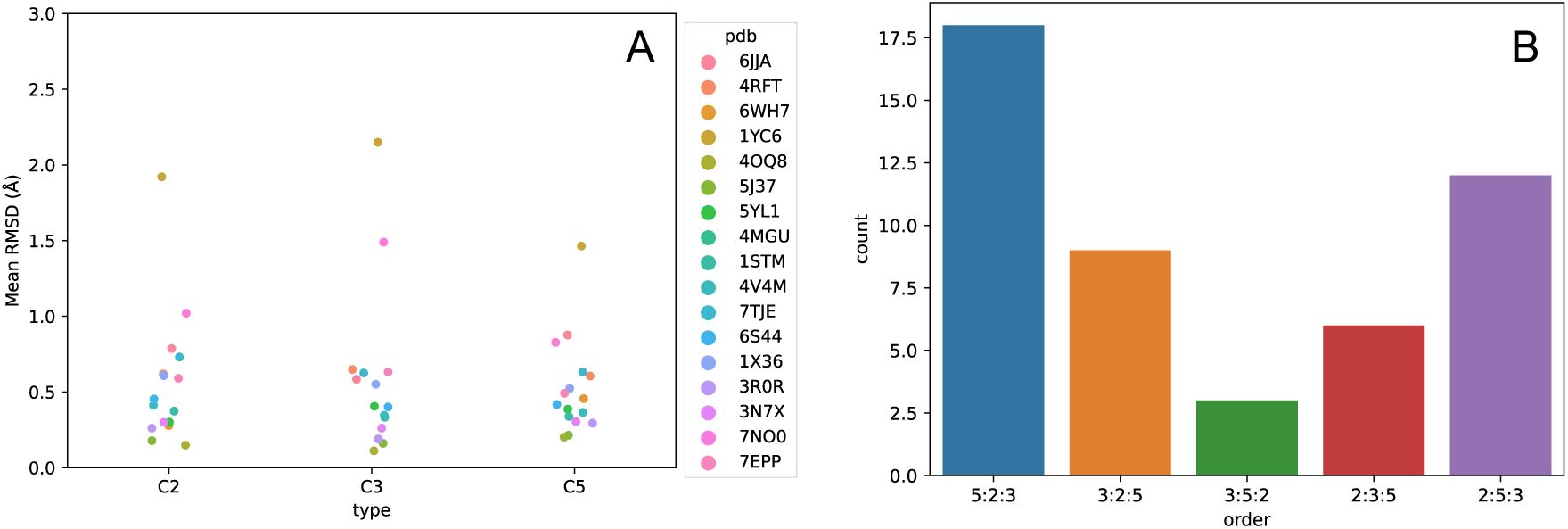
Diversity of oligomer structures in low-energy assembly models. A) The mean RMSD values over a set of the 2% lowest energy models for the dimer, trimer, and pentamer components of the capsid compared to the experimental structures. RMSD values were calculated over two consecutive protomers in the I:C2, I:C3 and I:C5 oligomers. Each T=1 has a different color in the plot. The plot was cut above 3 Å, excluding one system with mean RMSD values around 10Å. B) Relative order of mean RMSD values in interfaces. 5:2:3 corresponds to a system where the mean RMSD value for I:C5 is higher than I:C2, and I:C2 higher than I:C3.

### Hierarchy of interfaces in viral capsid

The relative size and stability of interfaces in the capsid shells can give insights into the most likely assembly pathway. The order of interface sizes was enumerated and counted, and the results are shown in Figure 7. The most common ranking of interfaces in virus capsids is having the largest interface in the trimer, followed by the pentamer (3:5:2). This is true also when considering stability, with the trimer being the most stable followed by the pentamer in most cases. Pentamers are commonly the biggest interface, but rarely the most stable in the T=1 set.

**Figure 7:**
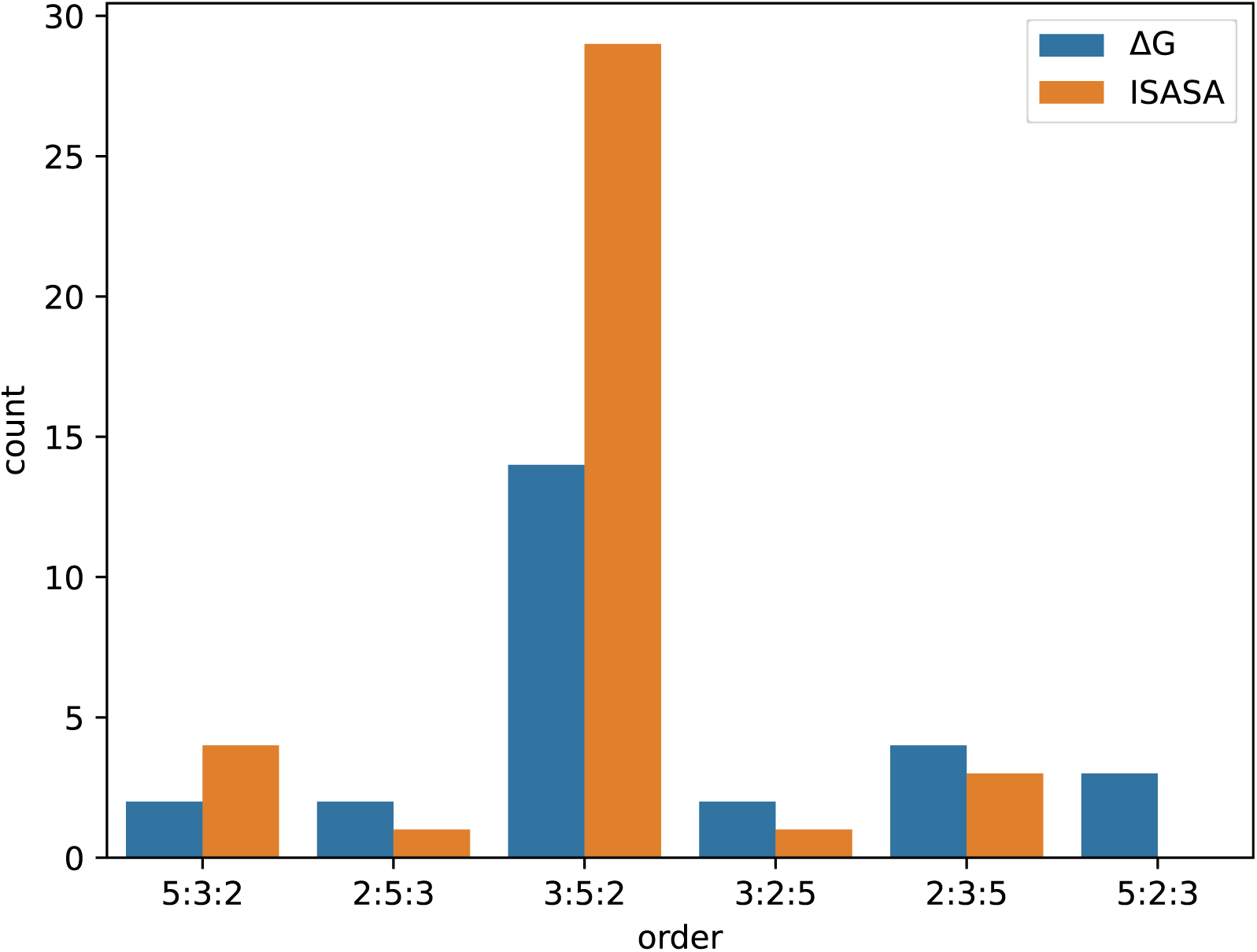
Relative size and stability of virus capsid interfaces. In 5:3:2 the 5-fold interfaces are bigger than the 3-fold and the 3-fold bigger than the 2-fold. For ΔG, the 5-fold is more stable than the 3-fold, etc. The y-axis shows the number of counts of each category. The ΔG data was filtered to remove systems with ΔG > 0, corresponding to non-converged energy refinements.

It is also interesting to determine the magnitude of the relative interface sizes and stabilities. Are capsids dominated by one big/stable interface with the other two interfaces being small and with limited stability? Or are the interface sizes and stabilities relatively evenly spread out between the interfaces? To answer that question, we calculated the relative size/stability of the second and third largest/most stable interface in each capsid. The results are presented in Figure 8.

**Figure 8:**
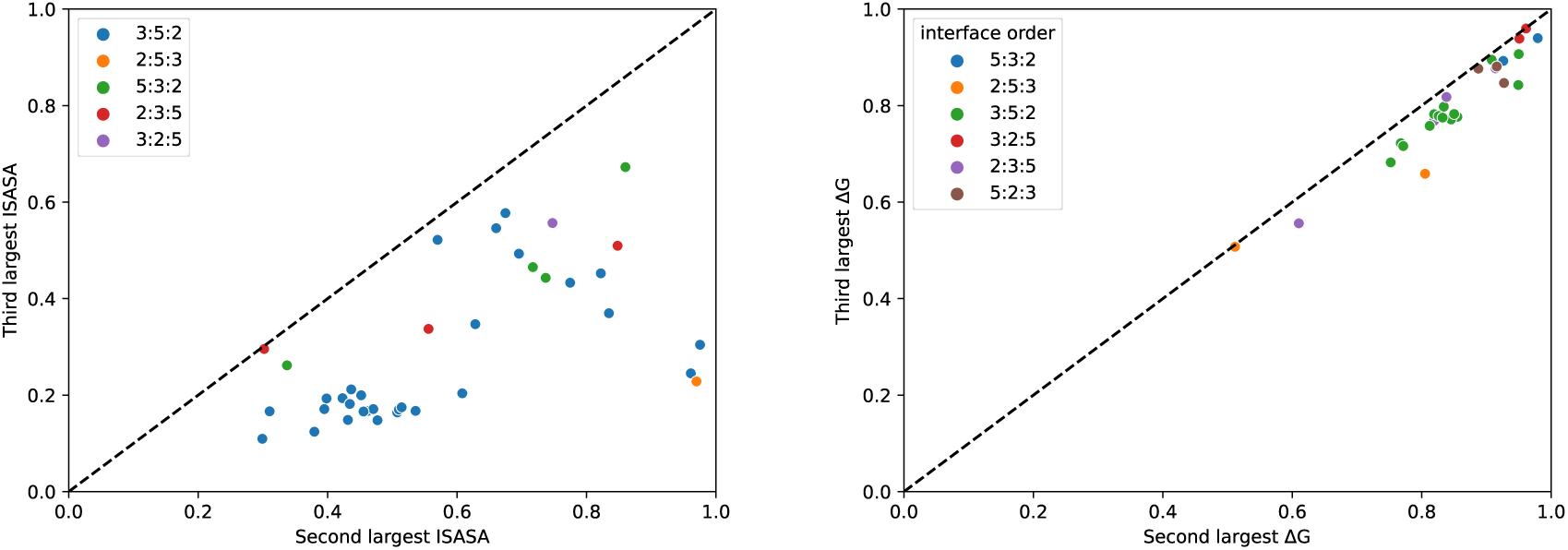
Relative size and stability of interfaces in virus T=1 capsid. Left, on the x-axis the interface size of the second largest interface divided by the largest interface size is shown. On the y-axis, the size of the third largest interface is divided by the largest interface size. Right, on the x-axis the ΔG of the second most stable assembly divided by the ΔG of the most stable interface is shown. On the y-axis, the ΔG of the third most stable interface is divided by the ΔG most stable assembly. The dotted line shows the diagonal, x=y.

If the capsid assembly is dominated by one big interface, and two smaller ones we expect the points to the lower left corner of the plot (Figure 8, left panel), whereas if all interfaces are of similar sizes, we expect points at the upper right corner. The third-ranked interface is typically below 50% of the size of the largest interface (mean 30% of the size of the largest), while the second-ranked interface is in the range 40-80% (mean 58%). Most assemblies therefore fall in the intermediate regime, where the interface size drops substantially from first-ranked to second, and second to third with a few examples where the second- and third-ranked are both small. Most of the 3:5:2, the most common type of relative order, have relatively small interfaces for the pentamer and dimer compared to the trimer.

The same analysis can be carried out with the computed ΔG-values per subunit by ranking them by order and dividing the second- and third-ranked values by the value of the best ΔG per subunit value (Figure 8, right panel). The points end up on the diagonal, suggesting that the second and third most stable interfaces are similar in their stability. The stabilities of the second and third are generally quite similar to the most stable one, with averages of 85 and 80% relative to the first-ranked interface, respectively.

## Discussion

The most parsimonious model for the evolution of complex assemblies like icosahedral virus capsids is that they have evolved from smaller oligomer building blocks. This is supported by analysis by Levy et al. demonstrating that cyclic and dihedral homomers typically conserve interactions in their subcomponent[20], such as dimeric complexes that form part of dihedral assemblies. They also looked at which type of interfaces were most likely conserved in the assemblies and found that larger interfaces were more likely conserved than small ones. They also experimentally measured the disassembly pathways in electrospray mass spectrometry and found a strong correlation with interface size, in which the biggest interfaces were the most stable ones. Applied to capsids, this would indicate that the oligomers with the largest interfaces are likely the evolutionary building blocks for capsids. In our set, this is typically trimers. Nonetheless, assembly pathways are not necessarily the reverse of disassembly pathways, and the disassembly pathway may be different in solution compared to the ones probed with electrospray mass spectrometry. The evolution of capsids is also likely more complex than for cyclic and dihedral assemblies. It is also important to note that the assembly process is not happening under equilibrium conditions. The self-assembly mechanism also depends on the relative rate of association and disassociation of different oligomeric species. Kinetics models of capsid models have been built on the assumption of similar association rates but with different dissociation rates for oligomers[7], something that is supported by experimental data correlating stability changes and off-rates in protein complex formation [38]. The energetic perspective presented in our analysis also highlights trimers as the most stable oligomers that may be building blocks for early capsid assembly. Nonetheless, there is a substantially lower difference in computed stability between dimers, trimers, and pentamers compared to the picture painted from the burial surface area. This would suggest that self-assembly may occur with a substantially more complex process than the simple model involving the initial formation of a single intermediate (such as trimers).

Experimental data and theoretical considerations have suggested that subunit interfaces are relatively weak in viral capsids[1, 12]. It would therefore be expected that computed stabilities and interface energetics would be more beneficial in homomeric oligomers compared to their viral counterparts. This was not observed in our data. The viral oligomers have fewer packing interactions, but this is counteracted by other interactions that result in similar computed interface stabilities. This apparent paradox can be resolved by considering factors that contribute to the free energy of binding but are not fully accounted for. First, the surface interior of viral capsids is highly positively charged, to interact with nucleic acid. This results in strong repulsive electrostatic interactions. The electrostatic cost of adding additional subunits to a growing capsid is not accounted for in calculations based on small oligomers. In addition, the model for long-range electrostatic in Rosetta energy function is over-simplified. Second, complex formation can be associated with conformational changes that are also not accounted for in our calculations. Smaller conformational changes upon binding can be modeled to some extent by refining the subunits in the unbound state as well. We do not find any substantial differences when this is done. However, this type of energy refinement can only model limited changes conformation. Third, there is a substantial component of rotational and translational entropy lost upon binding[39], a loss that increases with the size of the assembly.

Taken together, we suggest that while viral interfaces are larger and have differences in amino acid propensities, they do not have lower-quality intermolecular interactions compared to the corresponding homomeric interfaces. Instead, the reduced stability of assembled capsids may be due to the electrostatic cost of the binding of subunits with similar charges, conformational changes upon interface assembly, or loss of rotational-translation entropy. The benefit of the electrostatic repulsion model (and to some extent the entropic model) is that initial assembly can be fast and efficient due to the quality of the protein-protein interfaces, while still allowing for lower stability of partially and fully assembled capsids. Further research will have to be done to test these hypotheses.

Efficient self-assembly of virus capsids has been associated with error correction, where misassembled defective capsid states can disassemble and rebuild[19]. Our atomistic docking results suggest that T=1 capsids have very funneled energy landscapes, with much higher energy associated with alternative interface geometries. This type of specificity could be beneficial in driving assembly formation towards the final capsid geometry that must involve very highly uniform interfaces for the full capsid to be formed. However, an important reason for the formation of deeply funneled landscapes is also the icosahedral symmetry of the capsids, which imposes strong constraints on the movement of individual subunits in the lattice. It is likely that the demands on forming the experimentally observed subunit interfaces are reduced earlier in the assembly process where fewer subunits have associated into oligomers.

## Methods

### Data analysis

All data analysis was done using custom scripts develop in Python3[40], using the package scipy for statistical calculations[41] and Seaborn for plotting[42].

### Collection of protein structures

A list of T1 capsid PDB entries was downloaded from ViperDB[5]. This list was cross-referenced with the Protein Data Bank[25] to only select entries containing 60 subunits with Icosahedral symmetry that had higher resolution than 4Å. PDBs that could not be symmetrized were removed. To reduce the structural redundancy the list was furthermore filtered based on 70% sequence identity using CD-hit as follows:

> *cd-hit -i <sequences> -d 0 -o <output file> -c 0.7 -n 5 -G 1 -g 1 -b 20 -l 10 -s 0.0 -aL 0.0 -aS 0.0 -T 10 -M 32000*

Finally, non-viral structures were removed which yielded a set of 45 viral T1 structures with sequence lengths between 132 and 735 residues per subunit.

Homomeric assemblies of C_2_, C_3,_ and C_5_ symmetry were harvested from the protein data bank[25], requesting a chain length between 50-750 residues, spanning the length of structures in the T=1 set. C_2_ and C_3_ symmetries were further limited to a maximum 3Å resolution and C_5_ to 4Å resolution and all symmetries were clustered at 30% sequence identity.

### Selection of homomeric set with similar length distribution to the viral set

The chain lengths of the T=1 capsid proteins were calculated using a script developed with biopython[43]. The distribution of chain lengths was summarized as a histogram. To select a subset of homomers with similar chain lengths to this target distribution, histograms were calculated for the subset and the Jensen-Shannon divergence was minimized. To maintain a larger number of members in the selected ensemble, a cost for removing a structure from the set was introduced, resulting in the following cost function to be optimized:

> Cost = JSD(target || sampled set) + (selected number of structures/total number of structures)/5

Where 5 is a scaling factor. The cost was minimized using simulated annealing, based on a simanneal Python module for simulated annealing (https://github.com/perrygeo/simanneal).

### Calculation of structural and energetic metrics

The Rosetta macromolecular modeling package[36], through the python interface pyrosetta[44] was used to calculate structural and energetic metrics. Before analysis, the structures were symmetrized by using the symmetry modeling machinery in Rosetta[45]. The viral n-folds were extracted from the T=1 capsids before analysis. Interface Solvent Accessible Surface Area (ISASA) was calculated by evaluating the SASA in the bound minus the unbound state (generated by translating subunits symmetrically 2000 Å apart). SASA calculations was carried out using the SasaCalc class in Rosetta. Interface residues were calculated in a series of steps. In the first step, all residues having their CB atoms within 10Å of any other CB atom on another chain are found. The residues in the subset are then considered to be a potential interface residue if either of the two conditions is true. The first condition is that any atom of the residue is within 5.5 Å of any other atom on another chain in the subset. For the second condition, two vectors are drawn. One from the CA-to-CB atom of one residue and a CB-to-CB atom between two residues in the subset on opposite chains. If the angle between these two vectors is below 75 degrees, the pair is considered interface residues. Finally, to designate them as interface residues the residue-level ISASA must be above 0. Before analysis, the energetics and stereochemistry were optimized by carrying out a symmetric energy refinement with Rosetta (Symmetric FastRelax) [23]. These refinements were carried out with or without coordinate constraints on the Ca positions in the subunits. For energetic metrics, the bound relaxed structure is compared to unbound states in which the subunits are translated 2000 Å away perpendicular to their rotational symmetry axis. The side chains of the unbound states were optimized before energy evaluation to calculate ΔG values, and changes in van der Waals, hydrogen bonding, and solvation energy. The standard Rosetta energy function was used for these calculations[24]. The data presented in the manuscript was filtered to remove systems with suboptimal energy refinements by taking out systems with ΔG above 0, which were failures in convergence of the symmetrical FastRelax protocol. Also, interfaces with less than 2 interface residues were removed. For normalization with interface size, interfaces with less than 10 residues were removed.

### Energy landscape calculations of viral capsids

Energy landscapes for T=1virus capsids were generated with a symmetric version of the EvoDOCK protein-protein docking program. The method is presented in detail in Jeppesen and André[35], while the underlying differential evolution method used in EvoDOCK is presented in Varela et. al[34]. The selected T=1 capsids were analyzed to generate a symmetry definition used to recreate a fully symmetric system (icosahedral) system inside EvoDOCK. These six degrees of freedom used to define the symmetry of the system were then randomly perturbed as described in ref[35] for local recapitulation but a larger parametric space was sampled by setting the parameter bounds to *ψ*=[−60, 60], *θ*=[−60, 60], *φ=*[−60, 60], *z=*[−10, 10] *x=*[−10, 10] and *λ=* [−36, 36]. The structure and energetics reoptimized in a local docking simulation guided by the Rosetta energy function. For each simulation 100 starting orientations were generated before and 50 generations in the genetic algorithm were employed to generate the final ensemble.

To select the structures for energy landscape analysis the 45 T=1 structures were clustered at 40% sequence identity with CD-hit as described earlier. Then the set was sorted based on sequence length and the 20 structures with the shortest sequences that could be symmetrized with Rosetta were selected.

## Acknowledgments

This work was supported by the European Research Council (ERC) under the European Union’s Horizon 2020 research and innovation program [771820]

## References

[1] Perlmutter JD, Hagan MF. Mechanisms of virus assembly. Annu Rev Phys Chem. 2015;66:217–39.

[2] Rossmann MG. Structure of viruses: a short history. Q Rev Biophys. 2013;46:133–80.

[3] Bahadur RP, Rodier F, Janin J. A dissection of the protein-protein interfaces in icosahedral virus capsids. J Mol Biol. 2007;367:574–90.

[4] Ban N, McPherson A. The structure of satellite panicum mosaic virus at 1.9 A resolution. Nat Struct Biol. 1995;2:882–90.

[5] Montiel-Garcia D, Santoyo-Rivera N, Ho P, Carrillo-Tripp M, Brooks CL, Johnson JE, et al. VIPERdb v3.0: a structure-based data analytics platform for viral capsids. Nucleic Acids Res. 2021;49:D809–D16.

[6] Perlmutter JD, Hagan MF. Mechanisms of Virus Assembly. Annu Rev Phys Chem. 2015;66:1–23.

[7] Zlotnick A, Johnson JM, Wingfield PW, Stahl SJ, Endres D. A theoretical model successfully identifies features of hepatitis B virus capsid assembly. Biochemistry. 1999;38:14644–52.

[8] Law-Hine D, Zeghal M, Bressanelli S, Constantin D, Tresset G. Identification of a major intermediate along the self-assembly pathway of an icosahedral viral capsid by using an analytical model of a spherical patch. Soft Matter. 2016;12:6728–36.

[9] Oliver RC, Potrzebowski W, Najibi SM, Pedersen MN, Arleth L, Mahmoudi N, et al. Assembly of Capsids from Hepatitis B Virus Core Protein Progresses through Highly Populated Intermediates in the Presence and Absence of RNA. ACS Nano. 2020;14:10226–38.

[10] Janin J, Bahadur RP, Chakrabarti P. Protein–protein interaction and quaternary structure. Q Rev Biophys. 2008;41:133–80.

[11] Mateu MG. Assembly, stability and dynamics of virus capsids. Arch Biochem Biophys. 2013;531:65–79.

[12] Zlotnick A. Are weak protein-protein interactions the general rule in capsid assembly? Virology. 2003;315:269–74.

[13] Wang JC-Y, Chen C, Rayaprolu V, Mukhopadhyay S, Zlotnick A. Self-Assembly of an Alphavirus Core-like Particle Is Distinguished by Strong Intersubunit Association Energy and Structural Defects. ACS Nano. 2015;9:8898–906.

[14] Whitesides GM, Boncheva M. Beyond molecules: self-assembly of mesoscopic and macroscopic components. Proc Natl Acad Sci U S A. 2002;99:4769–74.

[15] Endres D, Zlotnick A. Model-based analysis of assembly kinetics for virus capsids or other spherical polymers. Biophys J. 2002;83:1217–30.

[16] Zlotnick A, Aldrich R, Johnson JM, Ceres P, Young MJ. Mechanism of capsid assembly for an icosahedral plant virus. Virology. 2000;277:450–6.

[17] Hagan MF, Chandler D. Dynamic pathways for viral capsid assembly. Biophys J. 2006;91:42–54.

[18] Schwartz R, Shor PW, Prevelige PE, Jr., Berger B. Local rules simulation of the kinetics of virus capsid self-assembly. Biophys J. 1998;75:2626–36.

[19] Lutomski CA, Lyktey NA, Zhao ZC, Pierson EE, Zlotnick A, Jarrold MF. Hepatitis B Virus Capsid Completion Occurs through Error Correction. J Am Chem Soc. 2017;139:16932–8.

[20] Levy ED, Boeri Erba E, Robinson CV, Teichmann SA. Assembly reflects evolution of protein complexes. Nature. 2008;453:1262–5.

[21] Cheng SS, Brooks CL. Protein-Protein Interfaces in Viral Capsids Are Structurally Unique. J Mol Biol. 2015;427:3613–24.

[22] Norn CH, André I. Computational design of protein self-assembly. Curr Opin Struct Biol. 2016;39:39–45.

[23] Tyka MD, Keedy DA, Andre I, DiMaio F, Song YF, Richardson DC, et al. Alternate States of Proteins Revealed by Detailed Energy Landscape Mapping. J Mol Biol. 2011;405:607–18.

[24] Park H, Bradley P, Greisen P, Liu Y, Mulligan VK, Kim DE, et al. Simultaneous Optimization of Biomolecular Energy Functions on Features from Small Molecules and Macromolecules. J Chem Theory Comput. 2016;12:6201–12.

[25] Berman HM, Westbrook J, Feng Z, Gilliland G, Bhat TN, Weissig H, et al. The Protein Data Bank. Nucleic Acids Res. 2000;28:235–42.

[26] Wilcoxon F. Individual Comparisons by Ranking Methods. Biometrics Bull. 1945;1:80–3.

[27] Bahadur RP, Chakrabarti P, Rodier F, Janin J. Dissecting subunit interfaces in homodimeric proteins. Proteins. 2003;53:708–19.

[28] Lawrence MC, Colman PM. Shape complementarity at protein/protein interfaces. J Mol Biol. 1993;234:946–50.

[29] Janin J, Bahadur RP, Chakrabarti P. Protein-protein interaction and quaternary structure. Q Rev Biophys. 2008;41:133–80.

[30] Chakravarty D, Guharoy M, Robert CH, Chakrabarti P, Janin J. Reassessing buried surface areas in protein-protein complexes. Protein Sci. 2013;22:1453–7.

[31] Kulshreshtha S, Chaudhary V, Goswami GK, Mathur N. Computational approaches for predicting mutant protein stability. J Comput Aided Mol Des. 2016;30:401–12.

[32] Valanciute A, Nygaard L, Zschach H, Maglegaard Jepsen M, Lindorff-Larsen K, Stein A. Accurate protein stability predictions from homology models. Comput Struct Biotechnol J. 2023;21:66–73.

[33] Chaudhury S, Berrondo M, Weitzner BD, Muthu P, Bergman H, Gray JJ. Benchmarking and analysis of protein docking performance in Rosetta v3.2. Plos One. 2011;6:e22477.

[34] Varela D, Karlin V, Andre I. A memetic algorithm enables efficient local and global all-atom protein-protein docking with backbone and side-chain flexibility. Structure. 2022;30:1550–8 e3.

[35] Jeppesen M, André I. Accurate prediction of protein assembly structure by combining AlphaFold and symmetrical docking. bioRxiv. 2023:2023.06.22.546069.

[36] Leman JK, Weitzner BD, Lewis SM, Adolf-Bryfogle J, Alam N, Alford RF, et al. Macromolecular modeling and design in Rosetta: recent methods and frameworks. Nat Methods. 2020;17:665–80.

[37] Ramisch S, Lizatovic R, Andre I. Exploring alternate states and oligomerization preferences of coiled-coils by de novo structure modeling. Proteins. 2015;83:235–47.

[38] Agius R, Torchala M, Moal IH, Fernandez-Recio J, Bates PA. Characterizing Changes in the Rate of Protein-Protein Dissociation upon Interface Mutation Using Hotspot Energy and Organization. Plos Comput Biol. 2013;9.

[39] Wand AJ, Sharp KA. Measuring Entropy in Molecular Recognition by Proteins. Annu Rev Biophys. 2018;47:41–61.

[40] Rossum GV, Drake FL. Python 3 Reference Manual. Scotts Valley, CA: CreateSpace.

[41] Virtanen P, Gommers R, Oliphant TE, Haberland M, Reddy T, Cournapeau D, et al. SciPy 1.0: Fundamental Algorithms for Scientific Computing in Python. Nature Methods. 2020;17:261–72.

[42] Waskom ML. seaborn: statistical data visualization. Journal of Open Source Software. 2021;6:3021.

[43] Cock PJ, Antao T, Chang JT, Chapman BA, Cox CJ, Dalke A, et al. Biopython: freely available Python tools for computational molecular biology and bioinformatics. Bioinformatics. 2009;25:1422–3.

[44] Chaudhury S, Lyskov S, Gray JJ. PyRosetta: a script-based interface for implementing molecular modeling algorithms using Rosetta. Bioinformatics. 2010;26:689–91.

[45] DiMaio F, Leaver-Fay A, Bradley P, Baker D, Andre I. Modeling Symmetric Macromolecular Structures in Rosetta3. Plos One. 2011;6.

