## Supplementary information for "Encoding of T=1 virus capsid structures through the interfaces of oligomer subcomponents"

### Supplementary Figures and tables

| PDB | Virus | Resolution (Å) | Reference |
|---|---|---|---|
| 6wh7 | Penaeid shrimp densovirus capsid | 2.78 | [1] |
| 8ep2 | Aleutian Mink Disease Virus capsid | 2.37 | [2] |
| 4g0r | H-1 Parvovirus capsid | 2.70 | [3] |
| 4cq8 | Bovine Parvovirus-1 capsid | 3.20 | [4] |
| 3r0r | Porcine Circovirus 2 capsid | 2.35 | [5] |
| 6lat | Hepatitis E virus capsid | 3.40 | [6] |
| 6bwx | Human Bufaviruses D capsid | 2.84 | [7] |
| 3n7x | Penaeus Stylirostris Densovirus capsid | 2.50 | [8] |
| 5j37 | beak and feather disease virus capsid | 2.30 | [9] |
| 5yl1 | Penaeus Vannamei Nodavirus capsid | 3.12 | [10] |
| 6x2k | Tusavirus 1 capsid | 2.88 | [11] |
| 4mgu | Acheta Domesticus Densovirus | 3.50 | [12] |
| 1dnv | Galleria Mellonella capsid | 3.60 | [13] |
| 7tje | Bacteriophage Q Beta Capsid | 2.80 | [14] |
| 7thr | Adeno-associated virus serotype 4 capsid | 2.21 | - |
| 1yc6 | Brome Mosaic Virus capsid | 2.90 | [15] |
| 7m3m | Canine Parvovirus capsid | 2.26 | [16] |
| 3p0s | Bombyx Mori Densovirus 1 capsid | 3.10 | [17] |
| 8ep9 | Human Parvovirus 4 capsid | 3.12 | [2] |
| 7epp | Alfalfa Mosaic Virus Coat Protein Virus-Like Particle | 2.40 | [18] |
| 7l0x | Human Bocavirus 2 capsid | 2.51 | [19] |
| 4oq8 | Satellite Tobacco Mosaic Virus | 1.45 | [20] |
| 6rpo | Bat Circovirus capsid | 2.39 | [21] |
| 1s58 | B19 Parvovirus capsid | 3.50 | [22] |
| 4rft | Grouper Nervous Necrosis Virus capsid | 3.10 | [23] |
| 1x9p | Human Adenovirus 2 Penton Base capsid | 3.30 | [24] |
| 7kp3 | Adeno-Associated Virus Serotype 5 capsid | 2.10 | [25] |
| 3ide | Aquabirnavirus capsid | 3.34 | [26] |
| 7u96 | Serpentine Adeno-Associated Virus capsid | 2.14 | [27] |
| 6jja | Giant Freshwater Prawn Macrobrachium Ro Extra Small Virus capsid | 2.91 | - |
| 1k3v | Porcine Parvovirus capsid | 3.50 | [28] |
| 1x36 | Sesbania Mosaic Virus capsid | 2.70 | [29] |
| 6x21 | Cutavirus capsid | 2.87 | [11] |
| 6wft | Bat Adeno-associat capsid | 3.03 | [30] |
| 7kfr | Adeno-Associated Virus capsid | 1.56 | [31] |
| 8er8 | Acheta Domesticus Segmented Densovirus capsid | 2.30 | - |
| 1dzl | L1 Protein of Human Papillomavirus 16 capsid | 3.50 | [32] |
| 6hcr | Adenovirus derived particle | 3.50 | [33] |
| 6b9q | Oncolytic Parvov Luiii capsid | 3.17 | [34] |
| 1stm | Satellite Panicum Mosaic Virus capsid | 1.90 | [35] |
| 7no0 | Mature Rous Sarcoma Virus capsid | 3.10 | [36] |
| 2gsy | infectious bursal disease virus capsid | 2.60 | [37] |
| 4v4m | Satellite Tobacco Necrosis Virus capsid | 1.45 | [38] |
| 6s44 | Faba Bean Necrotic Stunt Virus capsid | 3.19 | [39] |
| 6ssj | Human Endogenous Retrovirus capsid | 2.75 | [40] |

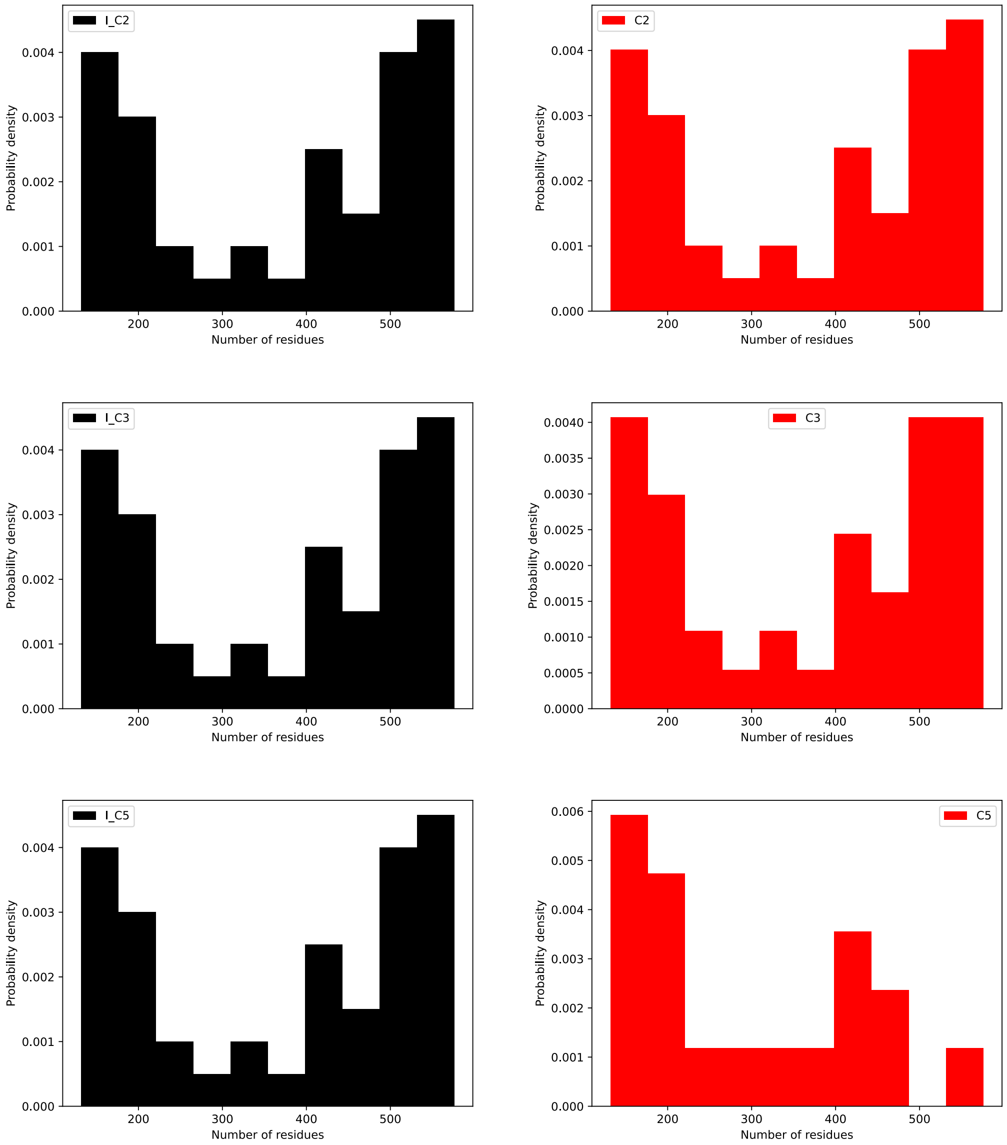


*Figure S1: Comparison of chain length distribution in capsid set (I:C2, I:C3, I:C5) and oligomer set (C2, C3 and C5 symmetric homomers) after ensemble selection to minimize the distance between distributions.*

### References

[1] Penzes JJ, Pham HT, Chipman P, Bhattacharya N, McKenna R, Agbandje-McKenna M, et al. Molecular biology and structure of a novel penaeid shrimp densovirus elucidate convergent parvoviral host capsid evolution. Proc Natl Acad Sci U S A. 2020;117:20211-22.

[2] Lakshmanan R, Mietzsch M, Jimenez Ybargollin A, Chipman P, Fu X, Qiu J, et al. Capsid Structure of Aleutian Mink Disease Virus and Human Parvovirus 4: New Faces in the Parvovirus Family Portrait. Viruses. 2022;14.
[3] Halder S, Nam HJ, Govindasamy L, Vogel M, Dinsart C, Salome N, et al. Structural characterization of H-1 parvovirus: comparison of infectious virions to empty capsids. J Virol. 2013;87:5128-40.
[4] Kailasan S, Halder S, Gurda B, Bladek H, Chipman PR, McKenna R, et al. Structure of an enteric pathogen, bovine parvovirus. J Virol. 2015;89:2603-14.
[5] Khayat R, Brunn N, Speir JA, Hardham JM, Ankenbauer RG, Schneemann A, et al. The 2.3-angstrom structure of porcine circovirus 2. J Virol. 2011;85:7856-62.
[6] Zheng Q, Jiang J, He M, Zheng Z, Yu H, Li T, et al. Viral neutralization by antibody-imposed physical disruption. Proc Natl Acad Sci U S A. 2019;116:26933-40.
[7] Ilyas M, Mietzsch M, Kailasan S, Vaisanen E, Luo M, Chipman P, et al. Atomic Resolution Structures of Human Bufaviruses Determined by Cryo-Electron Microscopy. Viruses. 2018;10.
[8] Kaufmann B, Bowman VD, Li Y, Szelei J, Waddell PJ, Tijssen P, et al. Structure of Penaeus stylirostris densovirus, a shrimp pathogen. J Virol. 2010;84:11289-96.
[9] Sarker S, Terron MC, Khandokar Y, Aragao D, Hardy JM, Radjainia M, et al. Structural insights into the assembly and regulation of distinct viral capsid complexes. Nat Commun. 2016;7:13014.
[10] Chen NC, Yoshimura M, Miyazaki N, Guan HH, Chuankhayan P, Lin CC, et al. The atomic structures of shrimp nodaviruses reveal new dimeric spike structures and particle polymorphism. Commun Biol. 2019;2:72.
[11] Mietzsch M, McKenna R, Vaisanen E, Yu JC, Ilyas M, Hull JA, et al. Structural Characterization of Cuta- and Tusavirus: Insight into Protoparvoviruses Capsid Morphology. Viruses. 2020;12.
[12] Meng G, Zhang X, Plevka P, Yu Q, Tijssen P, Rossmann MG. The structure and host entry of an invertebrate parvovirus. J Virol. 2013;87:12523-30.
[13] Simpson AA, Chipman PR, Baker TS, Tijssen P, Rossmann MG. The structure of an insect parvovirus (Galleria mellonella densovirus) at 3.7 A resolution. Structure. 1998;6:1355-67.
[14] Sungsuwan S, Wu X, Shaw V, Kavunja H, McFall-Boegeman H, Rashidijahanabad Z, et al. Structure Guided Design of Bacteriophage Qbeta Mutants as Next Generation Carriers for Conjugate Vaccines. ACS Chem Biol. 2022;17:3047-58.
[15] Larson SB, Lucas RW, McPherson A. Crystallographic structure of the T=1 particle of brome mosaic virus. J Mol Biol. 2005;346:815-31.
[16] Goetschius DJ, Hartmann SR, Organtini LJ, Callaway H, Huang K, Bator CM, et al. High-resolution asymmetric structure of a Fab-virus complex reveals overlap with the receptor binding site. Proc Natl Acad Sci U S A. 2021;118.
[17] Kaufmann B, El-Far M, Plevka P, Bowman VD, Li Y, Tijssen P, et al. Structure of Bombyx mori densovirus 1, a silkworm pathogen. J Virol. 2011;85:4691-7.
[18] Jeong H, Park Y, Song S, Min K, Woo JS, Lee YH, et al. Characterization of alfalfa mosaic virus capsid protein using Cryo-EM. Biochem Biophys Res Commun. 2021;559:161-7.
[19] Luo M, Mietzsch M, Chipman P, Song K, Xu C, Spear J, et al. pH-Induced Conformational Changes of Human Bocavirus Capsids. J Virol. 2021;95.
[20] Larson SB, Day JS, McPherson A. Satellite tobacco mosaic virus refined to 1.4 A resolution. Acta Crystallogr D Biol Crystallogr. 2014;70:2316-30.

[21] Nath BK, Das S, Roby JA, Sarker S, Luque D, Raidal SR, et al. Structural Perspectives of Beak and Feather Disease Virus and Porcine Circovirus Proteins. Viral Immunol. 2021;34:49-59.
[22] Kaufmann B, Simpson AA, Rossmann MG. The structure of human parvovirus B19. Proc Natl Acad Sci U S A. 2004;101:11628-33.
[23] Chen NC, Yoshimura M, Guan HH, Wang TY, Misumi Y, Lin CC, et al. Crystal Structures of a Piscine Betanodavirus: Mechanisms of Capsid Assembly and Viral Infection. PLoS Pathog. 2015;11:e1005203.
[24] Zubieta C, Schoehn G, Chroboczek J, Cusack S. The structure of the human adenovirus 2 penton. Mol Cell. 2005;17:121-35.
[25] Silveria MA, Large EE, Zane GM, White TA, Chapman MS. The Structure of an AAV5-AAVR Complex at 2.5 A Resolution: Implications for Cellular Entry and Immune Neutralization of AAV Gene Therapy Vectors. Viruses. 2020;12.
[26] Coulibaly F, Chevalier C, Delmas B, Rey FA. Crystal structure of an Aquabirnavirus particle: insights into antigenic diversity and virulence determinism. J Virol. 2010;84:1792-9.
[27] Mietzsch M, Hull JA, Makal VE, Jimenez Ybargollin A, Yu JC, McKissock K, et al. Characterization of the Serpentine Adeno-Associated Virus (SAAV) Capsid Structure: Receptor Interactions and Antigenicity. J Virol. 2022;96:e0033522.
[28] Wery JP, Reddy VS, Hosur MV, Johnson JE. The refined three-dimensional structure of an insect virus at 2.8 A resolution. J Mol Biol. 1994;235:565-86.
[29] Sangita V, Satheshkumar PS, Savithri HS, Murthy MR. Structure of a mutant T=1 capsid of Sesbania mosaic virus: role of water molecules in capsid architecture and integrity. Acta Crystallogr D Biol Crystallogr. 2005;61:1406-12.
[30] Mietzsch M, Li Y, Kurian J, Smith JK, Chipman P, McKenna R, et al. Structural characterization of a bat Adeno-associated virus capsid. J Struct Biol. 2020;211:107547.
[31] Xie Q, Yoshioka CK, Chapman MS. Adeno-Associated Virus (AAV-DJ)-Cryo-EM Structure at 1.56 A Resolution. Viruses. 2020;12.
[32] Chen XS, Garcea RL, Goldberg I, Casini G, Harrison SC. Structure of small virus-like particles assembled from the L1 protein of human papillomavirus 16. Mol Cell. 2000;5:557-67.
[33] Vragniau C, Bufton JC, Garzoni F, Stermann E, Rabi F, Terrat C, et al. Synthetic self-assembling ADDomer platform for highly efficient vaccination by genetically encoded multiepitope display. Sci Adv. 2019;5:eaaw2853.
[34] Pittman N, Misseldine A, Geilen L, Halder S, Smith JK, Kurian J, et al. Atomic Resolution Structure of the Oncolytic Parvovirus LuIII by Electron Microscopy and 3D Image Reconstruction. Viruses. 2017;9.
[35] Ban N, McPherson A. The structure of satellite panicum mosaic virus at 1.9 A resolution. Nat Struct Biol. 1995;2:882-90.
[36] Obr M, Ricana CL, Nikulin N, Feathers JR, Klanschnig M, Thader A, et al. Structure of the mature Rous sarcoma virus lattice reveals a role for IP6 in the formation of the capsid hexamer. Nat Commun. 2021;12:3226.
[37] Garriga D, Querol-Audi J, Abaitua F, Saugar I, Pous J, Verdaguer N, et al. The 2.6-Angstrom structure of infectious bursal disease virus-derived T=1 particles reveals new stabilizing elements of the virus capsid. J Virol. 2006;80:6895-905.
[38] Lane SW, Dennis CA, Lane CL, Trinh CH, Rizkallah PJ, Stockley PG, et al. Construction and crystal structure of recombinant STNV capsids. J Mol Biol. 2011;413:41-50.

[39] Trapani S, Bhat EA, Yvon M, Lai-Kee-Him J, Hoh F, Vernerey MS, et al. Structure-guided mutagenesis of the capsid protein indicates that a nanovirus requires assembled viral particles for systemic infection. PLoS Pathog. 2023;19:e1011086.
[40] Acton O, Grant T, Nicastro G, Ball NJ, Goldstone DC, Robertson LE, et al. Structural basis for Fullerene geometry in a human endogenous retrovirus capsid. Nat Commun. 2019;10:5822.
